# Cryo-EM structures reveal a dynamic transformation process of human alpha-2-macroglobulin working as a protease inhibitor

**DOI:** 10.1101/2022.05.10.491262

**Authors:** Xiaoxing Huang, Youwang Wang, Cong Yu, Hui Zhang, Qiang Ru, Xinxin Li, Kai Song, Min Zhou, Ping Zhu

**Author notes:** Corresponding author: Ping Zhu.

## Abstract

Human alpha-2-macroglobulin is a well-known proteases inhibitor against a broad spectrum of proteases. It also plays important roles in immunity, inflammation, and infections. Here, we report cryo-EM structures of human alpha-2-macroglobulin of the native state, the transformed state induced by its authentic substrate, human trypsin, and serial intermediate states between the native and the fully induced state. These structures exhibit distinct conformations, which reveal a dynamic transformation process of alpha-2-macroglobulin acting as a protease inhibitor. The results shed light on the molecular mechanism of alpha-2-macroglobulin entrapping substrates, and help to understand how alpha-2-macroglobulin possesses variant physiological functions.

## Introduction

As one of the most versatile proteins in body, human Alpha-2-macroglobulin (A2M) possesses multifaceted functions and plays important roles in variant physiological processes such as inflammation, immunity and infections (1, 2). As a well-known broad proteases inhibitor, A2M usually adopts two conformations in vivo, i.e., native and induced forms. Native A2M captures almost all classes of proteases (serine, carboxyl, thiol, metallo-) and structurally converts to induced A2M when it carries the preys to the target cells and get them swallowed. Both the native and induced forms of A2M bind a number of cytokines, growth factors to different degrees and with different efficiency to regulate their activities. Interestingly, induced A2M was found to perform more functions like activating the signaling cascades in target cell that facilitates neutrophil migration, improves cell division of macrophages, promotes bacterial phagocytosis and antigen presentation by macrophages (1, 3–10). Moreover, induced A2M was found to inhibit amyloid formation and hypochlorite-induced A2M dimer with potent chaperone activity helps to remove the damaged proteins through lipoprotein receptors (1, 10). Actually, the physiological roles of both native and induced A2M have been gradually being discovered presenting extremely sophisticated functions. The malfunction of A2M were found to associate with various major human diseases, such as Alzheimer’s disease, diabetes, and arthritis, cancers, and cardiovascular disease (1, 11–18).

Since the two major existing forms, i.e, native and induced forms, of human A2M *in vivo* serve as the basis to participate in all sorts of physiological processes, understanding the three-dimensional structures of A2M in both forms and how A2M converts from native to induced form are critical to reveal how hA2M perform its physiological functions. Previously, human A2M in a tetrameric form was reported to inhibit the substrate proteases via a mechanism called “Venus flytrap” which illustrates the main process of A2M transformation from native A2M to the induced A2M (2, 19). Several elements of human A2M were found critical in native A2M during the substrates trapping process, including a highly flexible bait region in the bait region domain (BRD) which can be recognized and hydrolyzed by different proteases(20–22), a highly reactive and hidden thioester site (CGEQ) in the thioester motif domain (TED) which is able to bind to the proteases covalently(23), and a cryptic receptor binding site containing key residues in the receptor binding domain (RBD) which will expose and bind to the receptor after the substrate protease captivity(24–26). The “Venus flytrap” is said to start with the proteolysis of the bait region by proteases, followed by a dramatic conformational change of A2M from the original state, i.e., native A2M (hereafter, nA2M), to an induced state, i.e., induced A2M (hereafter, iA2M), during the process the buried thioester site is exposed to bind the substrates instantly and covalently to prevent them from escaping. The RBD is then exposed to recognize its receptor, i.e., low-density lipoprotein receptor-related protein 1 (LRP1), on the target cells (27–29). The interaction of iA2M with its receptor leads to the clearance of A2M-substrate complex from circulation through cell endocytosis (1, 2). Alternatively, iA2M can be a diverse modulator as LRP1 potentially activates various signaling pathway, involving cell proliferation and adhesion, cytoskeleton reconstruction, and apoptosis in target cells (10), In addition, a number of cytokines and growth factors interact with nA2M but increasingly or preferentially with iA2M to prevent from proteolysis or to modulate their activities.

Structures of the native hA2M have been studied and obtained in low resolution by Cryo-EM or by the combination of negative staining transmission electron microscopy (NSTEM) and small-angle X-ray scattering (SAXS) (30, 31). However, the structure of induced hA2M trapped with authentic proteases have not been reported until now except for a proxy of which the substrate protease was replaced by a small nucleophile named methylamine (MA) which was found causes similar A2M conformational change with the real transformation but without any hydrolysis at the bait region (35–39). Structures of A2M in other eukaryotes such as *Xenopus laevis* have been reported recently (32). Although the structures of A2M in prokaryotes have been reported in high resolution compared with eukaryotes’, e.g., the ones of native and induced A2M in *Escherichia coli* (ECAM) (33, 34) or in *Salmonella enterica ser* (SCAM) (35), prokaryotic A2M is organized as monomer *in vivo* which are significantly different from eukaryotes’ tetramer. Taken together, despite the structures obtained above, the high resolution structures of human A2M (hereafter, hA2M) in both its native and authentic substrate protease induced states, particularly the structural information of those core elements (i.e., thioester bonds, bait region and RBD) are still necessary to be determined.

Here, we report a series of cryo-EM structures of hA2M, including the original native state (nA2M), the fully induced state by an authentic substrate, trypsin (hereafter, iA2M-trypsin), and a series of intermediates states between nA2M and iA2M. These structures revealed a whole process of A2M transformation from native to induced form upon hunting proteases. Our results shed light on the molecular features of A2M functioning as a broad proteases inhibitor and provide fundamental structural basis for understanding the intricate roles of A2M in a host of important physiological activities.

## Results

### Structures of native and induced A2Ms are distinct

Weighting about 180KD, human A2M consists of 11 individual domains, which include seven macroglobulin-like domains (MG1-MG7), a bait region domain (BRD) inserted in MG6, a complement C1r/C1s, Uegf, Bmp1 domain (CUB), a thioester motif domain (TED) inserted into the CUB domain, and a receptor binding domain (RBD) (Fig 1A).

**Fig.1.**
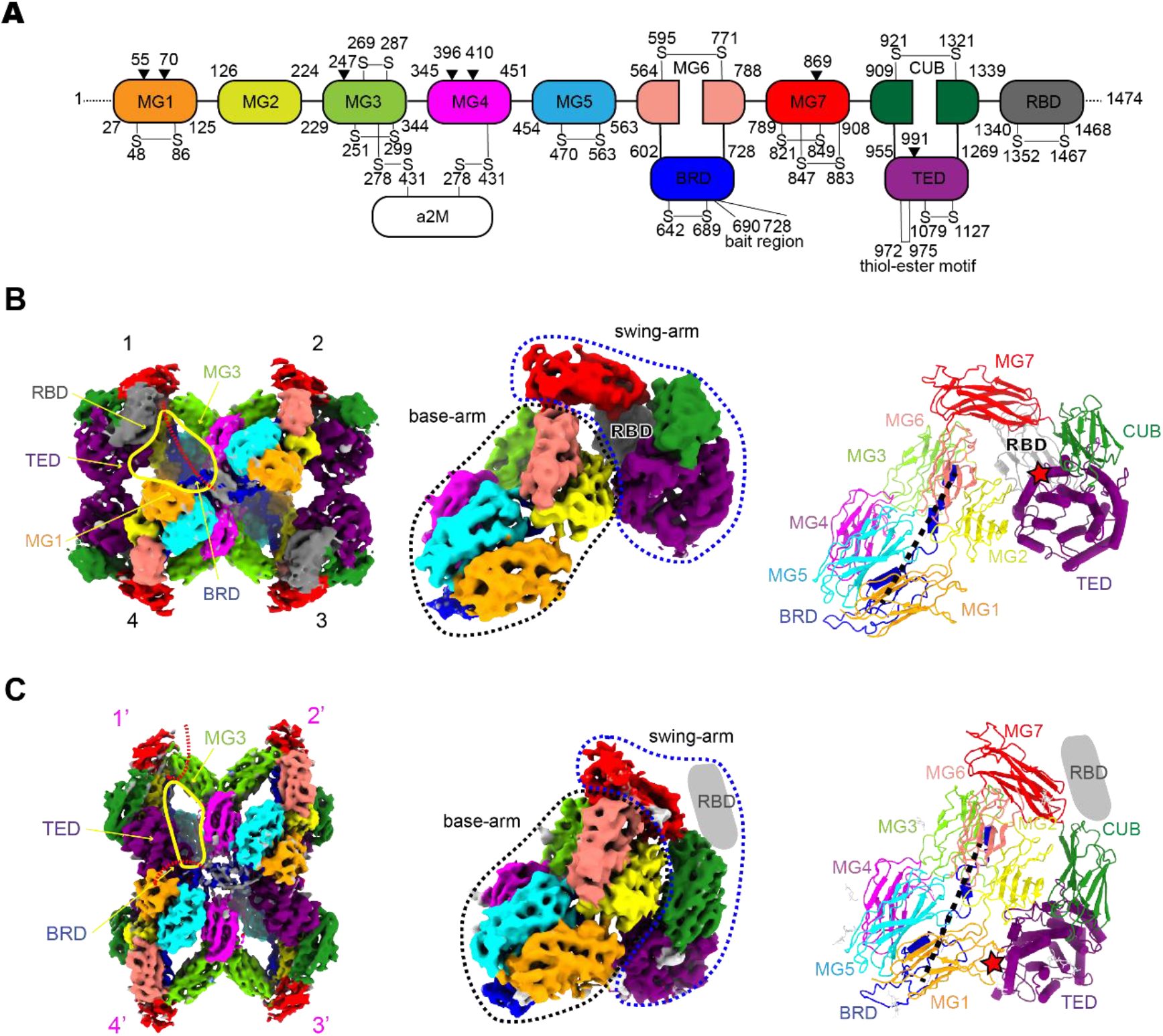
Structures of nA2M and iA2M are distinct. **(A**) Domain arrangement of human A2M. MG1-MG7, macroglobulin-type domains 1 to 7; BRD, bait-region domain; CUB, complement C1r/C1s, Uegf, Bmp1 domain; TED, thioester domain; RBD, receptor-binding domain. Black triangles pinpoint the glycosylation sites. **(B-C)** the density map for tetramer (left), the density map for monomer (middle), and atomic coordinate for monomer (right) of nA2M (B) and iA2M (C). The domains are colored as in (A). The black dotted lines in the coordinate panels represent the unresolved part of bait region. The red stars indicate the locations of thioester bond.

We purified human A2M (hA2M) from blood plasma (*SI Appendix*, Fig S1) and determined the overall structure of native A2M, i.e., nA2M, through cryo-EM to 5.3 Å resolution (Fig 1B, left, Fig S2), and pushed the resolution of native monomer to 3.9 Å by combining all datasets and locally refining focused on native monomer, eventually built an atomic structure model (Fig 1B, middle and right, Fig S2). To acquire a high resolution structure of human A2M induced by its authentic substrates, we treated the isolated hA2M, which possessed proteases entrapping activity (*SI Appendix*, Fig S1C), with trypsin in a 1:2 molar ratio to achieve the induced form of A2M, i.e., iA2M-trypsin. In line with the previous studies (23, 36), our isothermal titration calorimetry (ITC) assay indicated that hA2M mixed with trypsins in 1:2 molar ratio lead to a binding ratio of 1:1.2 (*SI Appendix*, Fig S3F), which is sufficient for producing fully induced architecture of A2M. 3D density map was reconstructed by cryo-EM single particle analysis at the resolution of 3.7 Å (Fig 1C, left, Fig S3). Comparing the overall architectures between nA2M and iA2M-trypsin (Fig 1B and C, left), the nA2M presents a global “extended” structure composed of four identical “extended” monomers, while iA2M-trypsin present a compact “retracted” structure composed of four identical “retracted” monomers, instead. A large cavity (about 48 Å in diameter) could only be observed in each monomer of nA2M (Fig 1B, left, yellow circle), which is big enough for the entry of the substrate proteinases. Upon the encapsulation of the substrate proteases, however, the cavity turns smaller (around 25 x 38 Å) (Fig 1C, left, yellow circle), which very likely prevents the captured substrates from escaping(37). Despite the overall configurations of nA2M (Fig 1B, left) and iA2M (Fig 1C, left) appear very different, the monomers both adopt a similar triangle-shaped hairpin conformation but with different vertex angles (Fig 1B and 1C, middle). The circular dichroism (CD) assays indicated that both nA2M and iA2M have a consistent pattern of secondary structures (*SI Appendix*, Fig S4), which suggests the sub-domain structures of A2M monomers are well preserved throughout the whole transformation. As shown in the structure of nA2M monomer (Fig 1B, middle), the first six macroglobulin-like domains, i.e., MG1-MG6 (MG ring), constitute one arm which is termed as the “base-arm”. Following MG6, MG7 connects the first arm to the second arm of the hairpin made of MG7-RBD, which is termed as the “swing-arm”. That is, MG7 serves as a hinge domain between the two arms. BRD, in which the cleavage of its core sequence, i.e., bait region, by substrate proteases triggers the first step of proteases capture (20), is inserted into the MG6 and spans about 85Å throughout the monomer from the bottom to top. The CUB domain, linked to MG7, forms a β-sandwich comprising two antiparallel four-stranded β-sheets. The TED domain containing six α-hairpins was found inserted into the outer surface of CUB, which embraces the critical highly reactive thioester motif in TED in a position to bind to proteinases covalently, a key for A2M functioning as the proteases inhibitor. The C-terminal domain of A2M, RBD, which is responsible for receptor binding, is readily visible in nA2M and binds to TED. Comparing the monomers of nA2M with iA2M, there’s no appreciable conformational change within the base-arm, while significant changes could be observed on the swing arm, which results in a huge structural difference of the monomer from the “extended” to “retracted” conformation (Fig 1B and 1C, right). Of these, the manner of the structural change located in the swing arm (including RBD) will be discussed below.

Although the bait region domain (BRD) is not well resolved, possibly owing to its high flexibility, the upstream and downstream of BRD in both nA2M and iA2M have been partially resolved (*SI Appendix*, Fig S5). In nA2M, part of flexible BRD extends inward in the entrance of the cavity (30), which likely serve as an entrance guard (Fig 1B left, red dashed line). The upstream and downstream of BRDs in nA2M and iA2M show noticeable different configurations (*SI Appendix*, Fig S5, Fig 1B and 1C, left), implying a conformational change induced by the proteolysis of trypsin in the bait region.

### Serial structures of A2Ms reveal a dynamic changing process of A2M transformation

Besides the structures of nA2M and iA2M-trypsin, we also resolved a series of hA2M purified from serum in other forms with multiple conformations at the resolutions from 5.3 to 8.2 Å respectively by 3D classification and refinement (*SI Appendix*, Fig S2), which represent variant intermediate states between nA2M and iA2M. All of these resolved structures taken together, vividly show a dynamically changing process of hA2M transforming from native to induced state (Fig 2A). Interestingly, although a series of hA2M structures with varied configurations were reconstructed, only two kinds of distinct conformations of hA2M monomers, i.e., “extended” (Fig 2A, gray) and “retracted” (Fig 2A, magenta), were found in the structures which build up all different forms of hA2M. In which, the “retracted” monomer is same with the one we resolved in iA2M-trypsin, while the “extended” monomer is same with the one in nA2M. Therefore, the different hA2M forms with changing ratio between “retracted” and “extended” monomers ranging of 0:4, 1:3, 2:2, 3:1and 4:0 (Fig 2A and *SI Appendix*, Fig S2C) can be explained as the dynamic transforming process with monomers “retracted” one by one from the initiate nA2M state with four “extracted” monomers.

**Fig.2.**
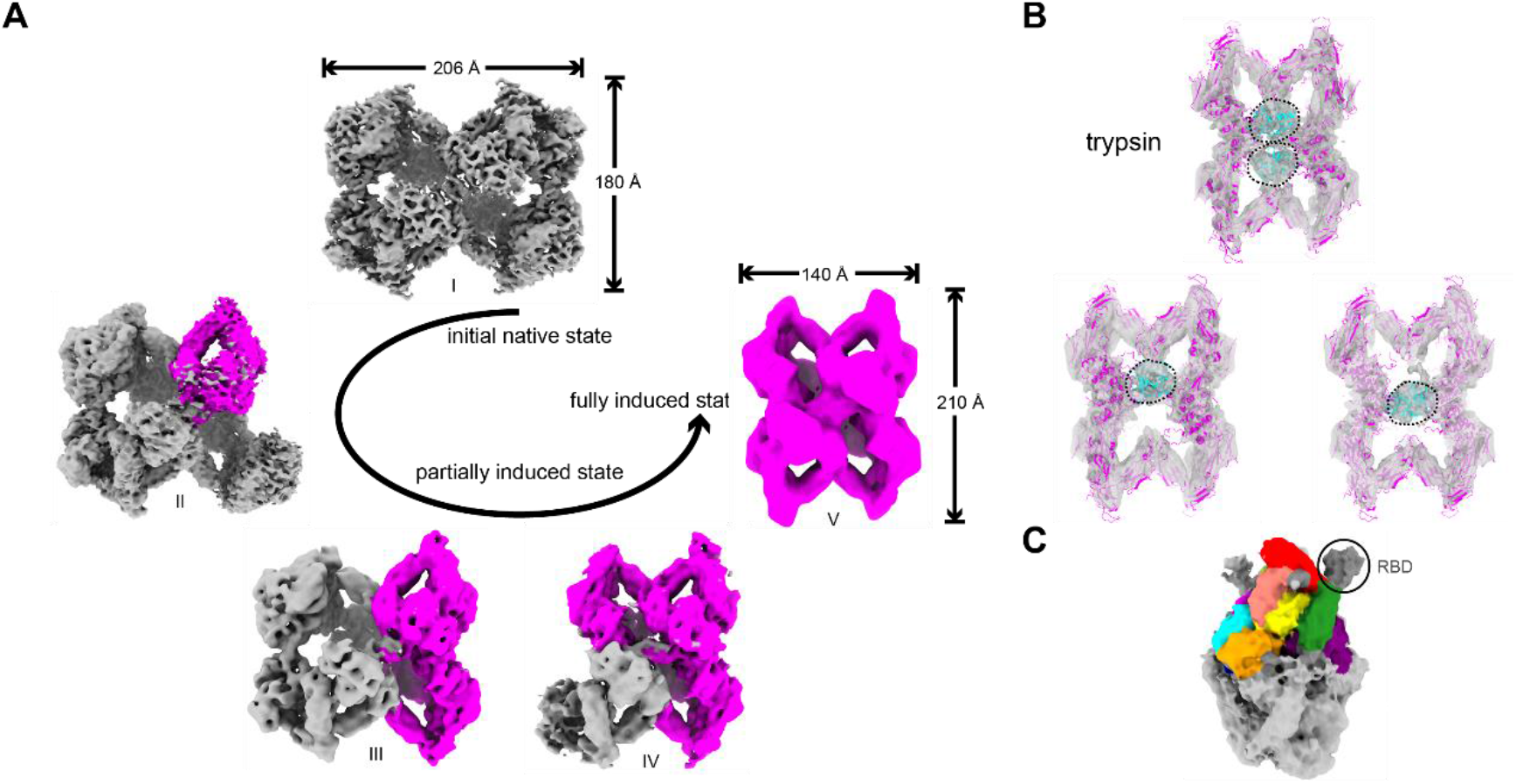
Cryo-EM structures of human A2M at different states and iA2M-trypsin entrapped one or two molecules of trypsin. **(A)** Cryo-EM structures of A2M purified from human blood at different states, including the native state (I), partially induced intermediate states (II, III, IV) and fully induced state (V). Two types of monomers are colored in gray (native) and magenta (induced), respectively. **(B)** Central slice of iA2M-trypsin density map fitted with coordinates of iA2M and trypsin (PDB: 1H4W, cyan) showing the densities of one or two trypsin molecules inside. **(C)** Reconstructed density of iA2M-trypsin displayed at a low contour level shows the flexible density of RBD (circled).

Interestingly, there is a minor group of hA2M with four “retracted” monomers isolated from human serum (Fig 2A, V), of which the overall structure is same with iA2M-trypsin but hard to observe the details in it due to poor resolution (8.2 Å). Comparatively, the structure of iA2M-trypsin provides more conformational details benefiting from its higher resolution (Fig. 1C). Although iA2M-trypsin, i.e., hA2M trapped with authentic proteases, trypsin, presents a similar overall architecture to that of iA2M-MA, A2M induced by MA, a small nucleophile (38), that is, a configuration 210 Å in length and 140 Å in width with four identical “retracted” monomers (Fig 1C), remarkable differences between the structures of iA2M-trypsin and iA2M-MA could be noticed. First, unlike the empty-caged structure of iA2M-MA, the density of substrate protease, i.e., trypsin could be clearly observed in the lumen of reconstructed maps of the iA2M-trypsin (Fig 2B). Interestingly, the reconstructed structures showed that the trypsin could be trapped by A2M with either two molecules in the lumen or only one in the lumen but in differential locations (Fig 2B). In consistent with it, the cross-linking mass spectrometry (XL-MS) indicated that the entrapped trypsin could be imprisoned within A2M in multiple orientations but with some preferred orientations and interactions (*SI Appendix*, Fig S6). Second, despite RBD is not well resolved in iA2M-trypsin, it was found in an orientation totally different from the one in iA2M-MA at low contour level (Fig 2C), indicating a conformational change of RBD induced by the authentic substrate protease, trypsin.

### Conformational change within or between monomers from nA2M to iA2M

From nA2M to iA2M, significant conformational changes, including the intra-monomeric configurations, and the inter-monomeric conformations, could be visibly identified (Fig 3). Within one monomer, there’s no appreciable change in the base-arm, while major changes could be observed on the swing arm, i.e. MG7, CUB, TED, RBD (Fig 3A). The most remarkable change was found between the CUB and TED domains, that is, TED rotates by 45° along with CUB in iA2M compared to that in nA2M (Fig 3B, left). As a consequence, the buried thioester site in nA2M bounces out and binds to the proteases in iA2M (Fig 2B). Besides this, the swing-arm swings toward the base-arm making the vertex angle between the MG6 and MG7 narrower by 30° (Fig 3B, middle), while the angle between the CUB and MG7 becomes 20° wider (Fig 3B, right). In contrast to sticking to the TED domain in nA2M (Fig 3A, left), the RBD domain stretches outward in iA2M-trypsin (Fig 3A, right and Fig 2C), which could interact with its receptor, e.g., LRP1, on the target cells. Taken together, the conformational changes in all of the above elements results in the monomer’s change from the “extended” to “retracted” conformation.

**Fig.3.**
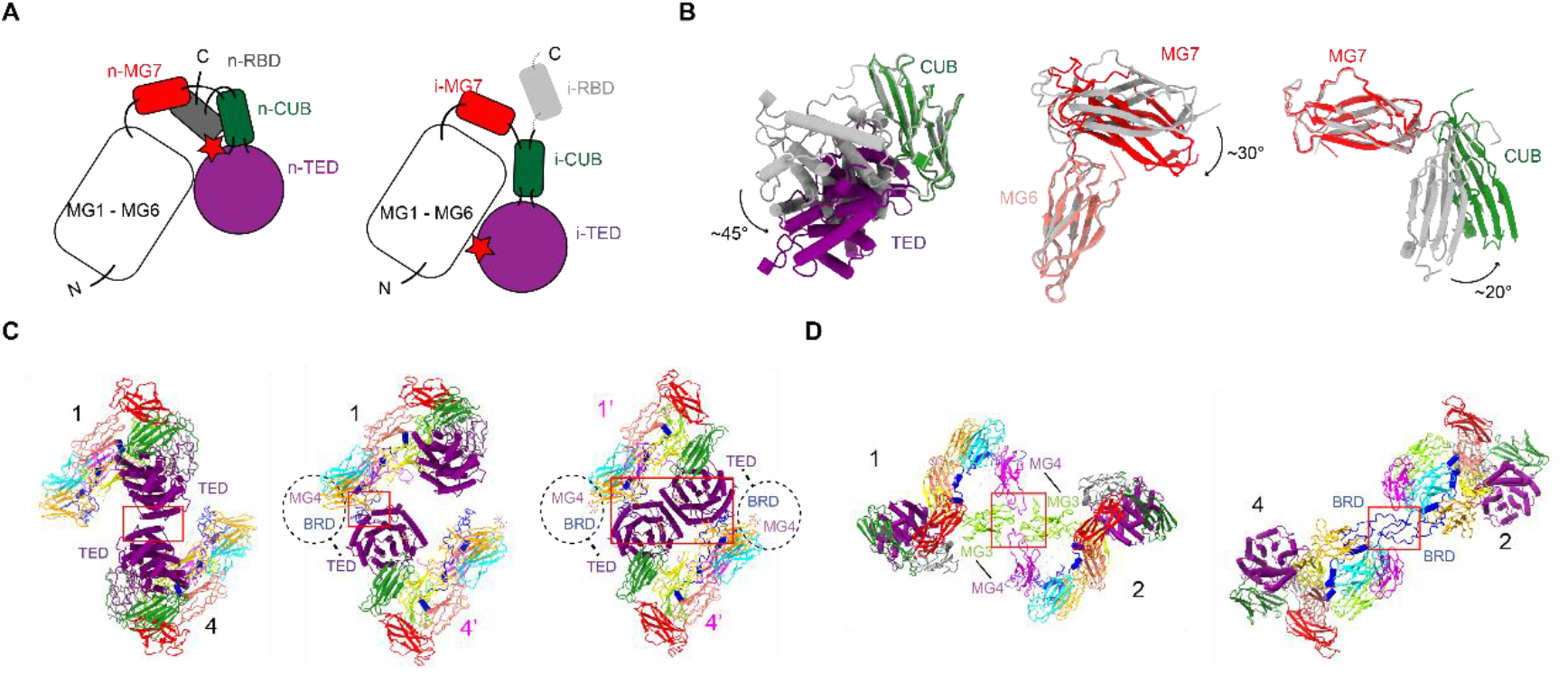
Conformational change within or between monomers from nA2M to iA2M. **(A)** Schematic domain arrangement of nA2M monomer (left) and iA2M monomer (right). **(B)** The structural alignments of CUB-TED (left), MG6-MG7 (middle) and MG7-CUB domains (right) between nA2M (gray) and iA2M (color). **(C)** Surface contacts between n-n monomers (left), n-i monomers (middle), and i-i monomers (right). **(D)** The consistent interaction of the disulfide bonds between two disulfide-linked monomers throughout A2M transformation (left). The consistent interaction of the “tethering loops” between two opposite monomers throughout A2M transformation. Magnified views and details of interactions are shown in Fig S7 (right).

In addition to the conformational changes among the sub-domains within one monomer, the overall architecture and the interfaces among monomers, i.e., monomer-monomer interactions, which also play important roles in the tetrameric assembly and transformation of A2M, are also subject to noteworthy changes during the process of A2M transformation. Firstly, the interactions between the two longitudinally vicinal monomers, i.e., 1-4 or 2-3, change dramatically during the transformation from nA2M to iA2M induced by substrates (Fig 3C). In nA2M, the two vicinal monomers, e.g., 1-4, in the native form (hereafter, n-monomers) interplay with each other through two α-helices in regions of R1073-F1126 of TED (Fig 3C, left and *SI Appendix*, Fig S7A). However, in the intermediate state, i.e., partially induced form of A2M, one of the two n-monomers, e.g., monomer 4, turns into the induced form (monomer 4’, hereafter i-monomer) while the other one, e.g., monomer 1, stays unchanged (Fig 3C, middle). Consequently, the interaction between two n-monomers, e.g., 1-4, in nA2M is broken, while new interactions between n-monomer (the H1020-V1120 region in TED) and i-monomer (the region of S66-V122 in MG4 and P633-C642 in BRD), e.g., 1-4’, are established (*SI Appendix*, Fig S7B). Notably, in the fully induced form, the two vicinal i-monomers, e.g., 1’-4’, present three interacted areas, i.e., the region of H1020-V1120 in TED of one i-monomer with that of S66-V122 in MG4 and P633-C642 in BRD upstream of the other i-monomer respectively, and the regions of I1005-R1014 in TED of both i-monomers contact to each other (Fig 3C right and *SI Appendix*, Fig S7C). Noticeably, the area of interactions between the two vicinal i-monomers increases significantly from 456 Å^2^ in the native form (Fig 3C, left), to 8609 Å^2^ in the induced form (Fig 3C, right), which significantly helps to hold the two disulfide-linked dimers together and keep the tetramer as a unit. The stronger interactions result in a much more stable structure of iA2M (Fig 1C, left) than that of nA2M (Fig 1B, left). This is in agreement with the previous study that the overall structure of iA2M would not be altered even all of the disulfide bonds were destroyed by dithiothreitol (DTT)(39).

To be worthy of note, two kinds of monomer-monomer interactions in nA2M, i.e., the disulfide linkage (C278 in MG3 with C431 in MG4 of the neighboring monomer) between two lateral monomers, i.e., 1-2 and 3-4 (Fig 3D left and *SI Appendix*, Fig S7D), and the interplay between tethering loops (C642-D665) of the two opposite monomers, i.e., 1-3 and 2-4 (Fig 3D right and *SI Appendix*, Fig S7E), were found persistent and unchanged throughout the A2M transformation process. In consistent with it, these two interactions were reported essential in the stabilization of A2M tetramer, as the reduction of the disulfide bonds between C278 of MG3 and C431 of MG4 by dithiothreitol (DTT) would cause the disassembly of nA2M into dimers or monomers (39), while the symmetrical interaction of tethering loops from the two opposite monomers seems the solely bondage to link double disulfide-linked dimers as a tetramer(30).

### Elements critical for the assembly of nA2M and proteases trapping

The hydrophobic pockets have been reported critical for protecting the highly reactive thioester from premature hydrolysis by proteases in A2M and other thioester containing proteins (40–44). However, it remains unclear how the “hydrophobic pocket” is constituted in human A2M. We built the structural model of human nA2M (Fig 4A) and found the “hydrophobic pocket” of human nA2M was constituted by a group of hydrophobic residues, e.g., M1378, Y1418, Y1452, Y1453 of RBD and M968, Y970, F1028 of TED, which is embraced between RBD and TED domains (Fig 4B and *SI Appendix*, Fig S8A). Interestingly, the hydrophobic residues of human A2M seems a combination of those from the prokaryotic members of thioester contained proteins e.g., ECAM, SCAM, and from the eukaryotic members, e.g., *Anopheles gambiae* TEP1, C3(41, 42), and presents a stronger hydrophobic environment than that in other species (*SI Appendix*, Fig S8). In this pocket, the TED domain appears like a “cup” that is covered by a “lid” (RBD), while the thioester bond formed by the side chains of C972 and Q975 is deeply buried in the “cup” and sheltered by the “lid” (Fig 4A). The cleft between the “cup” and “lid” appears narrow, making the sheltered thioester bond only accessible for small molecules, e.g., MA, but not big substrate proteases(39). Moreover, the cavity for the substrate protease entrance in nA2M is blocked by the bait region, which only allows small molecules like MA but not bigger ones like proteases to slip into and interact with the thioester bond (Fig 1B, left) (39). These results suggest the cleavage of the bait region, which shelters the cavity for the entrance of substrate protease, initiates a structural conformation change in nA2M. This conformational change leads to the exposure and subsequent attack of thioester bond by the substrate proteases, and induces a further conformational change of A2M to the fully induced state, i.e., iA2M. Therefore, although the attack of thioester bond by MA without bait region proteolysis may destroy the hydrophobic interactions between TED and RBD, thus converse nA2M into a similar form with iA2M(45–47), the cleavage of bait region, other than the attack on the thioester bond, appears the initiating step to induce the structural change of nA2M *in vivo*, as the thioester is not accessible for substrate proteases without the conformational change.

**Fig. 4.**
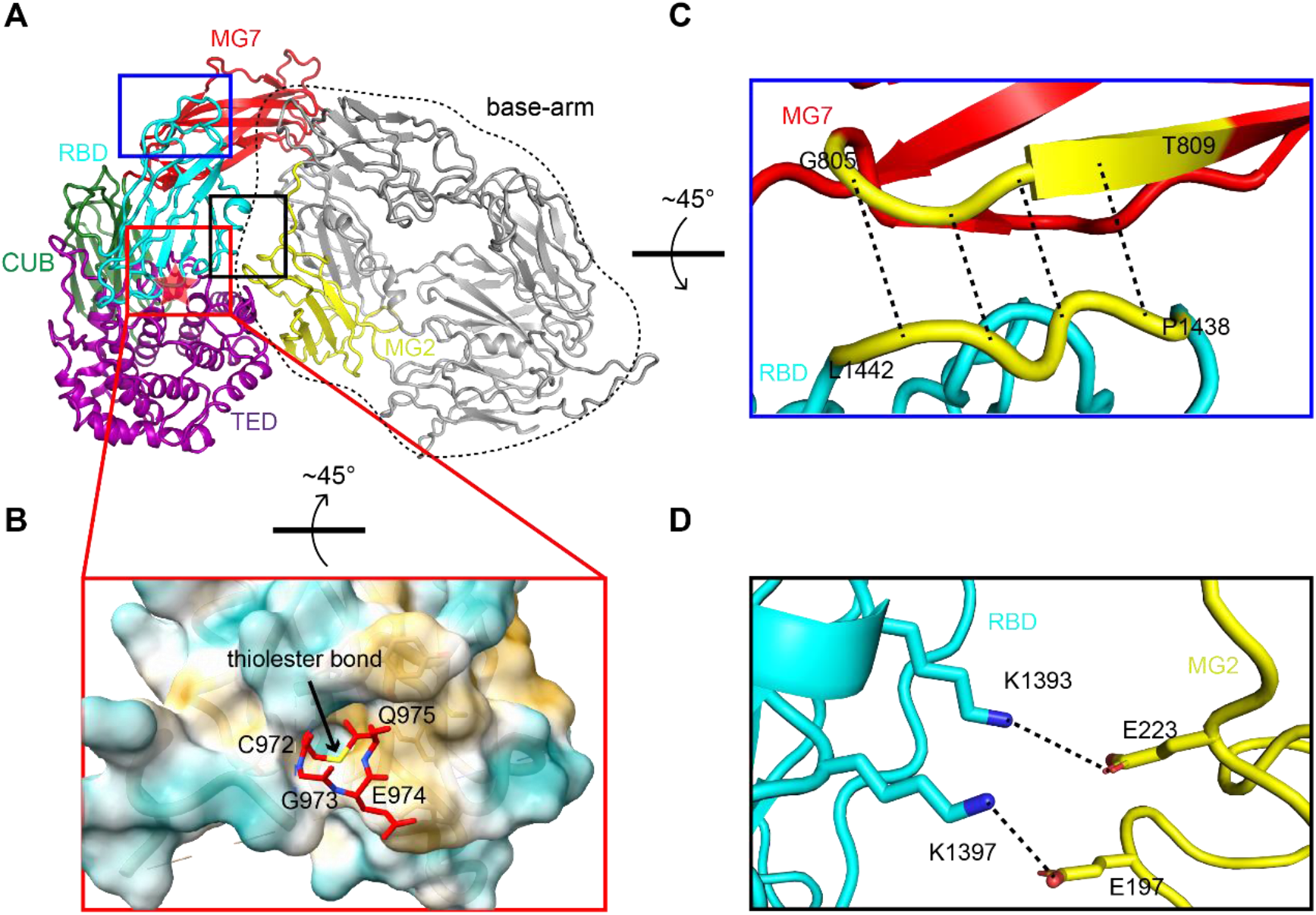
Hydrophobic pocket and receptor binding domain (RBD) in nA2M. **(A)** The overall configuration of nA2M monomer shown in ribbon mode. The domains of the swing-arm, i.e., TED, RBD, MG7, and CUB, and MG2 domain of the base arm are colored and labeled. Location of the hidden thioester bond is indicated by a transparent red star. **(B)** Zoom view of the red box in (A) shows the hydrophobic pocket protecting thioester bond from hydrolysis. The CGEQ residues in the thioester site are shown in red while the thioester bond is shown in yellow. Residues contributing to the formation of the hydrophobic pocket are shown in orange (see Fig S8*a* for details). **(C)** Zoom view of the blue box in (A) shows the interactions between RBD and MG7 helping to stabilize RBD in nA2M. **(D)** Zoom view of the black box in (A) shows the hidden residues K1393 and K1397 of RBD and their potential interactions with E197 and E223 in MG2.

The receptor binding domain (RBD) plays a key role for iA2M to bind to its receptor, e.g., LRP1, which initiates the endocytosis of target cell to intake iA2M-proteases complex to be digested. In our structures, the RBD was resolved well in nA2M (Fig 1B), but only partially in iA2M (Fig 2C), suggesting a more stable configuration of RBD in the context of nA2M. In nA2M, the interaction between RBD and TED plays an important role to stabilize the RBD (Fig 4A). Apart from it, MG7 also participates in the stabilization, in which the adjacent sheets of MG7 and RBD are within the distance of main chain hydrogen bonds and form an anti-parallel configuration (Fig 4C). Meanwhile, two lysine residues, i.e., K1393 and K1397, which were previously reported critical for the binding of RBD to LRP1(48), are found hidden in a cavity encircled by RBD, MG7, and MG3, and interact with E223 and E197 of MG2 in an electrostatic interaction manner (Fig 4D). Consistently, the mutations of these two lysine residues were reported to largely abolish the ability of RBD binding to LRP1(48). These two concealed lysine residues, therefore, protect nA2M from premature binding to LRP1 without carrying any cargo. Nevertheless, upon the substrates engagement, interfaces between RBD and TED, MG7, MG2 respectively which are presented in nA2M (Fig 4A), are vanished. The hidden residues K1393 and K1397 in iA2M are exposed to bind LRP1 on the target cells. This is in agreement with the previous report that only iA2M carried with proteinases can be quickly endocytosized by cells but not empty nA2M(45).

### A working model of the substrate protease inhibition process by human A2M

Our structures of hA2M in different forms, including the native form (nA2M), induced form (iA2M), and sequential intermediates during transformation, revealed a continuous changing process of A2M entrapping proteases. Based on these structures and previous studies, a working model of hA2M inhibiting substrate proteases is proposed (Fig 5).

**Fig.5.**
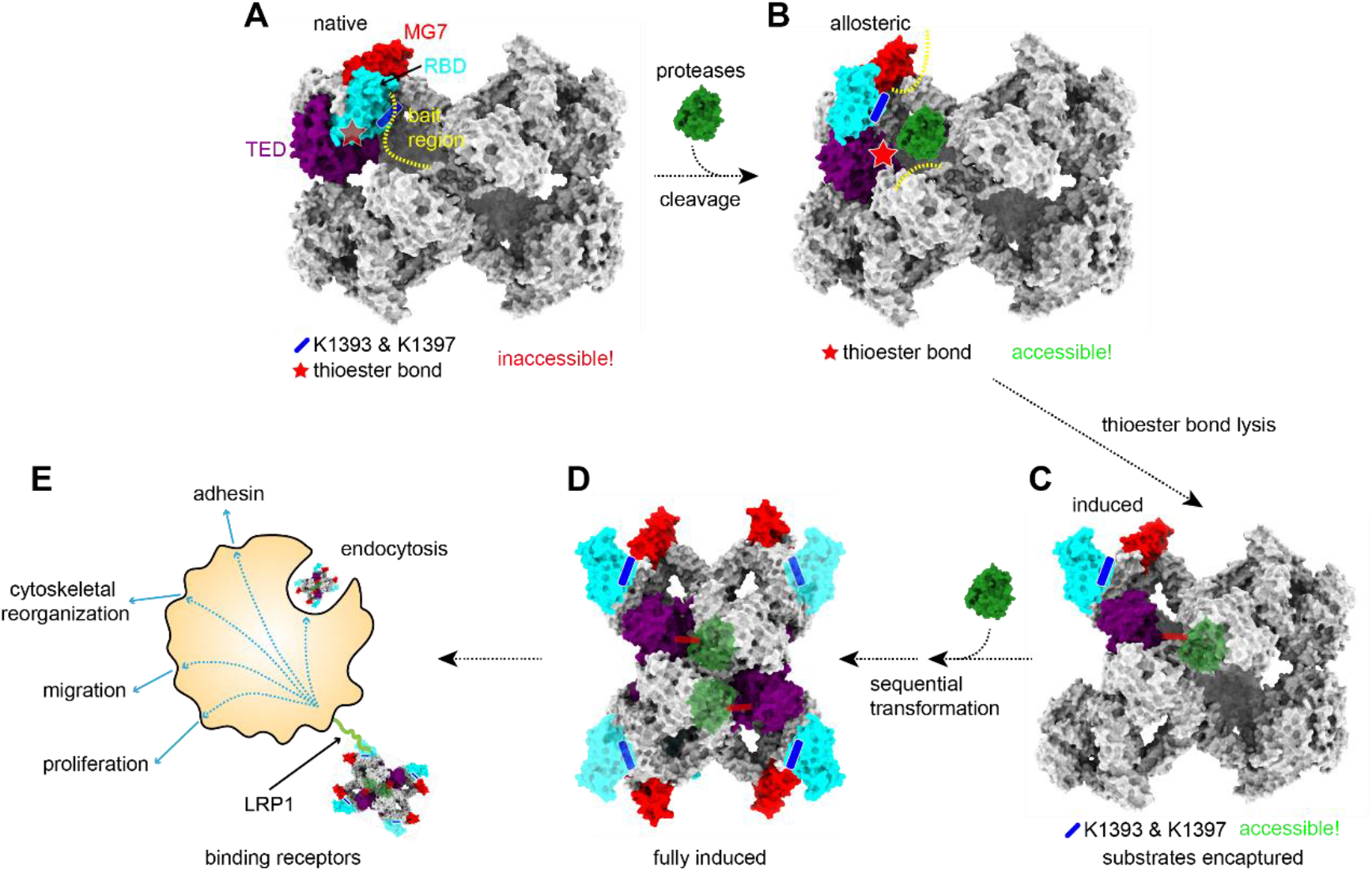
A working model of human A2M as proteases inhibitor. **(A)** nA2M is constituted by four “extended” n-monomers, in each of which the bait region (yellow dotted line) blocks the entrance of proteases. The interactions between RBD (cyan) and TED (purple)/MG7 (red) stabilize the RBD, build the “hydrophobic pocket” to embrace the thioester bond (transparent red star) and make the thioester and receptor binding residues (K1393 and K1397, transparent blue stick) inaccessible for proteases and receptors (LRP1), respectively. **(B)** The cleavage of bait region by proteases makes the n-monomer allosteric and the thioester site spatially exposed to the substrates, while RBD is not fully liberated. **(C)** The attack on the thioester site by proteases transfers the allosteric monomer to the fully induced i-monomer, in which the two lysine residues, K1393 and K1397 of RBD are exposed, and the thioester bond covalently binds the substrate protease. **(D)** Fully induced iA2M with four “retracted” monomers and encaptured proteases after sequential monomers transformation. **(E)** iA2M binds to LRP1 and initiates the endocytosis of iA2M-cargo complexes by target cell, which could alternatively modulate cell migration, proliferation, phagocytosis, etc., depending on the circumstances.

The native form of hA2M, i.e., nA2M, assembles as a fully “extended” conformation with four monomers “stretch out”, in which, both the thioester bond protected by the “hydrophobic pocket” and the key lysine residues, K1393 and K1397, in RBD are hidden and inaccessible for protease and LPR1 receptor respectively. Meanwhile, the large cavity for proteases entry is blocked by the flexible bait region of BRD (Fig 5A). When A2M is attacked by proteases, the hydrolysis of the bait region by substrate proteases causes a conformational change of the corresponding monomer in nA2M to an allosteric state which destroys the “hydrophobic pocket” and partially exposes the thioester site (Fig 5B). Then the highly active thioester bond becomes reachable and is attacked by protease which causes a further conformational change to convert the allosteric A2M monomer into the fully induced state, i.e., i-monomer, at the same time, a covalent bond between the opened thioester of i-monomer and substrate is formed and the hidden lysine residues for receptor binding expose to be accessible for LRP1 (Fig 5C). The transition from the n-monomer to i-monomer is then transferred to its neighbors, making the remaining n-monomers convert to i-monomers sequentially with or without more substrate involved, finally shapes a fully induced A2M with four monomers “retracted”, i.e., iA2M (Fig 5D). As a result, the flexible exposed RBD of iA2M can be recognized by LRP1 receptor and binds to the target cells, which initiates the endocytosis of the iA2M-cargo complexes, eliminate them from circulation, and perform other enormous physiological functions (Fig 5E).

## Discussion

Human alpha macroglobulin, typified by hA2M, constitutes as much as 8-10% of total serum proteins. HA2M participates in a variety of important physiological processes such as facilitating cell migration and proliferation, modulating leukocytes, binding damaged proteins, inhibiting the proteases secreted by invading microorganism etc., invariably based on its unique feature of being a universal protease inhibitor (10, 49). In order to interpret the mechanism of hA2M inhibiting proteases and its biological relevance, we determined a series of cryo-EM structures of human alpha-2-macroglobulin including nA2M, iA2M and the sequential intermediates between the two states. These structures collectively reveal a continuous conformational change of hA2M transformation from native to induced form during the whole process of entrapping substrate proteases.

Based on the structures, a model for protease entrapment by hA2M was proposed. In nA2M, the thioester is buried between RBD and TED and surrounded by a strong hydrophobic pocket, which protects it from the attack of substrate proteases. The receptor binding domain (RBD), which protects the thioester site, and prevents the key lysine residues for receptor binding from premature releasing, is stabilized by TED, MG7, and MG2. In addition, the flexible bait region in BRD acts as an entrance guard to block the entry of substrate protease. Although the attack by the small molecule, e.g., MA, without cleaving the bait region, can also transfer nA2M to a similar induced form, the proteolysis of bait region by protease seems a necessary initial step for the conformational change from nA2M to iA2M *in vivo*, as the thioester site is not reachable for substrate protease before its exposure. After the proteolysis of bait region, sub-domains in the swing arm of nA2M, i.e., MG7, CUB, TED, and RBD, are rearranged, which breaks up the “hydrophobic pocket” and drives the TED domain to shift towards the protease and bind it covalently. The RBD domain unwraps and exposes the critical lysine residues to its receptor. As a consequence, the entirely transformed iA2M specifically binds to its receptor LRP1 and is internalized by the target cell. This is the way A2M gets rid of the excessive proteases in blood. Alternatively, the receptor LRP1 has the potential to activate various signaling pathway in target cells, including cell proliferation, cytoskeleton reconstruction, cell adhesion and apoptosis(10), which enable A2M to play an important role to modulate multiple further physiological processes, involving neutrophils migration and adhesion, macrophages phagocytosis, etc.

Different from “Venus trap”, nA2M in prokaryotes such as ECAM and SCAM seems involve a different way to convert into iA2M when trapping substrate proteases, that’s called “Snap-trap”. That was indicated by the phenomenon in which prokaryotic nA2M would not transform by MA interaction instead of eukaryotic ones, but would definitely transfer into iA2M when cleaved by substrate proteases (33–35). Lack of necessary structural information of BRD, we cannot provide the exact explanation for the difference. But the recent work in which recombinant hA2M was mutant with modified BRD to perform XL-MS indicates that BRD should orient inwards and can be accessible from inside A2M (30). Agreed with this outcome and previous analysis (33), the BRD can be much shorter in prokaryotic A2M (about 25 residues) than in eukaryotic A2M (about 39 residues). The short BRD could greatly restrict the movement of RBD which is crucial to keep the whole structure stable. Instead, long BRD could allow a wide range of movement of RBD even if BRD remains intact. This may partially explain why MA involvement without BRD cleavage could lead to hA2M transformation while it’s not the case in ECAM. It appears that the interaction of MA with thiolester in hA2M created some instability to “hydrophobic pocket”, which somehow opens the latter and transformation happens. The reason behind this needs further investigation.

Interestingly, among various forms of hA2M we have obtained, there is a kind of induced form only contains one molecule of trypsin, while its four monomers all exhibit “retracted” conformation. This could mean the structural change of one monomer triggered by the proteases proteolysis is somehow transmitted to other uncleaved monomers to form four “retracted” i-monomers, which is in agreement with the previous report that, in the condition of A2M excess, the stoichiometry is 4:1 between cleaved thioester and trypsin added, while with two cleaved BRD (23). In line with this, the intermediates we solved also exhibit sequentially “retracted” monomers. These findings further revealed the details of hA2M transformation. However, through what manner, the conformational change can be transmitted within the A2M tetramer remains unexplored.

Taken together, we solved a seris of structures of hA2M in both native and induced form, and the intermediates between the two as well. These structures collectively exhibit the whole process of hA2M transformation upon proteases entrapment. The critical structural elements involved in hA2M inhibition are analyzed. These results shed light on the mechanism of hA2M broadly inhibiting proteases and provide structural clues for hA2M participating in many other physiological functions.

## Materials and Methods

Human A2M was purified and assayed for proteases entrapment activity through standard techniques. I-A2M was produced by treating A2M with trypsins. The structures of proteins were studied by Cryo-EM. A detailed description can be found in the *SI Appendix*, which also includes supplementary figures and table.

## Acknowledgments

We thank Jianhui Li, Zhenwei Yang, Yuanyuan Chen, Xiaoxia Yu at the Core Facility for Protein Research (Institute of Biophysics, Chinese Academy of Sciences) for assistance with CD and ITC experiments. We thank Gang Ji, Xiaojun Huang, Boling Zhu, Xujing Li at the Center for Biological Imaging (CBI), Core Facility for Protein Sciences, Chinese Academy of Sciences for assistance with EM data collection. We thank Jifeng Wang, Zhensheng Xie, Xiang Ding, Mengmeng Zhang (Proteomics, Institute of Biophysics, Chinese Academy of Sciences) for assistance with the Q-Exactive mass spectrometer analysis.

## Abbreviations

A2M: alpha-2-macroglobulin
cryo-EM: cryo-electron microscopy
hA2M: human alpha-2-macroglobulin
nA2M: native alpha-2-macroglobulin
iA2M: induced alpha-2-macroglobulin
iA2M-trypsin: A2M induced by trypsin
MA: methylamine
iA2M-MA: A2M induced by MA
n-monomer: native monomer
i-monomer: induced monomer
XL-MS: cross-linking mass spectrometry
NSTEM: negative staining transmission electron microscopy
SAXS: small-angle X-ray scattering
BRD: bait region domain
TED: thioester motif domain
RBD: receptor binding domain
MG1-MG7: macroglobulin-type domains 1 to 7
CUB: complement C1r/C1s, Uegf, Bmp1 domain
ITC: isothermal titration calorimetry
CD: the circular dichroism
3D: three dimentional
LRP1: low-density lipoprotein receptor-related protein 1
ECAM: A2M in *Escherichia coli*
SCAM: *Salmonella enterica ser*

## Declarations

### Ethics approval

Not applicable

### Consent to participate

Not applicable

### Consent for publication

Not applicable

### Availability of data and material

Cryo-EM reconstruction density maps of iA2M-trypsin and one form of purified A2M (1r3e) have been deposited into the Electron Microscopy Data Bank with access codes EMD-32052 and EMD-32051, while the coordinates of them have been deposited into the Protein Data Bank with access codes 7VOO and 7VON, respectively. Other forms of purified A2M (0r4e, 2r2e, 3r1e, 4r0e) have been deposited in the Electron Microscopy Data Bank with access codes EMD-32053, EMD-32054, EMD-32055, EMD-32057, respectively.

### Code availability

Not applicable

### Author Contributions

P.Z. and X.H. conceived the research. X.H., C.Y. and Q.R. prepared samples with assistance from X.L. and K.S.. X.H. and Y.W. collected the cryo-EM data with the assistance from M.Z. Y.W. and H.Z. processed the cryo-EM data. H.X., Y.W. and P.Z wrote the manuscript with inputs from all authors.

