## Supplementary information for "Cryo-EM structures reveal a dynamic transformation process of human alpha-2-macroglobulin working as a protease inhibitor"

**Experimental Procedures**

**Protein purification.** Human native A2M was isolated from blood plasma as previously described (1). Briefly, human blood from healthy consenting donors was centrifuged at 1000g, 4℃ for 30 min to pellet cells. The plasma was then loaded on a HiTrap chelating column (GE Healthcare) for further purification. The bound proteins were eluted with 500 mM imidazole, 0.5 M NaCl, 20 mM HEPES, pH 7.2, and dialyzed against phosphate buffered saline (PBS; 137 mM NaCl, 2.7 mM KCl, 1.7 mM KH2PO4, 10 mM Na2HPO4, pH 7.4). The dialyzed proteins were fractionated by a HiPrep 26/60 Sephacryl S-300 gel filtration column and the A2M containing fractions were pooled and stored at 4°C. The iA2M-trypsin was produced by treating purified A2M with human trypsin (Yaxin) at an 1:2 molar ratio, 37℃ for 1hr, in PBS.

**Native polyacrylamide gel electrophoresis (PAGE).** Electrophoretic mobility of A2M or iA2M-trypsin was examined by native PAGE using Native PAGE^TM^ 4–16% Bis-Tris gels (Life Technologies), and Native PAGE^TM^ running buffer (50mM BisTris, 50mM Tricine, pH 6.8) (1), according to the manufacturer’s instructions.

**Protease entrapment activity of alpha-2-macroglobulin.** The proteases entrapment activity of A2M was measured using Nα-Benzoyl-L-arginine ethyl ester (BAEE) as the substrate of trypsin, following the previous studies with a little modification (2). Briefly, 1mM BAEE (Alfa Aesar) was prepared in BAEE buffer (67mM NaH2PO4, pH 7.6). A2M (100μg) and varying amounts of trypsin (1-11μg) were incubated in total 0.1 ml 1mM BAEE buffer at room temperature (RT) for 10 minutes, before soybean trypsin inhibitor (STI, AMRESCO) was added to the solution. After a further incubation (10 min, RT) with 30μg STI, 1.4ml 1mM BAEE was added to the solution and the conversion of BAEE to Nα-Benzoyl-L-arginine (BA) at RT was measured at 253 nm using a UNICO 2800UV/VIS SPECTROPHOTOMETER. Trypsin activity was denoted by increased absorption per minute.

[**Isothermal titration calorimetry**](https://www.baidu.com/link?url=w0aIQG_89L8KGzbYLv-4o7zs6O0eOmn8QfOSnVeICpemw4D_eWqrSwTG_7kf5cc1zNu5rPhdK9DxumUbGFcrpufqXUdQfXeuV-k5qqrfeZzDh3oQIE4c8sX6F8ndhJoof3dUotdk266EOkgzYpVLs_&wd=&eqid=e49982d400051c01000000035d5a8917) **(ITC).** ITC measurements were carried out with MicroCalorimeter ITC200 (Microcal LLC) at 25°C. All of the proteins were dissolved in PBS (pH 7.4). The titration processes were performed by injecting 20× 2 µl aliquots of trypsin sample in a syringe (concentration of 42 µM) into stirred A2M sample (concentration of 2 µM) in the calorimeter cell at time intervals of 120 s. A background experiment using the buffer alone was performed to correct for the heats of mixing and dilution. Curve fitting was undertaken in ORIGIN 8.0 and fitted by the one-site-binding model.

**Circular dichroism (CD) spectroscopy.** CD experiments were performed using Chirascan plus (Applied Photophysics). A2M (0.22mg/ml) in PBS (pH 7.4) was analyzed using a 0.1 cm path-length cuvette. For secondary structure analysis, five spectra of each protein sample and phosphate buffer were recorded between 200 and 260 nm at 20°C (using a scan speed of 50 nm/min with a 1 nm bandwidth and a 4 s response time). The spectra of the samples were averaged and corrected for the signal generated by the buffer alone.

**Cryo-EM data acquisition.** For iA2M-trypsin, a 4 μL volume of the trypsin-treated A2M was applied to a glow-discharged Quantifoil copper grid and vitrified by plunge freezing in liquid ethane using a Vitrobot Mark (Thermo Fisher) with blotting time of 3s. Data collection was performed on a Titan Krios microscope (Thermo Fisher) operated at 300kV and equipped with a field emission gun, a Gatan GIF Quantum energy filter and a Gatan K2 Summit direct electron camera (Gatan) in super-resolution mode. The calibrated magnification was 130,000× in EFTEM mode, corresponding to a pixel size of 1.04 Å. The automated software SerialEM was used to collect 1,600 movies at a defocus range between 1.8 and 2.5 μm. Each exposure (10s exposure time) comprised 32 sub-frames amounting to a total dose of 60 electrons Å^-2^ s^-1^. nA2M cryogenic sample preparation and data acquisition were performed similarly as above, with 5,400 movies collected.

**Cryo-EM image processing.** Micrograph movie stacks were corrected for beam-induced motion using MotionCor2 (3). The contrast transfer function parameters for each dose weighting image were determined with Gctf (4). Particles were initially auto picked with Gautomatch (<https://www.mrc-lmb.cam.ac.uk/kzhang/>) without template and extracted with a 270-pixel by 270-pixel box for iA2M and nA2M datasets. Reference-free 2D-class average was performed using RELION (5), and the well-resolved 2D averages were subjected to another iteration particle auto picking as a template with Gautomatch. After iterative 2D-class average in RELION, particles with the best-resolved 2D averages were selected for initial model generation and 3D classification using RELION. The classes with similar features were merged for further auto-refinement with a sphere mask, and post-processed with 3-pixel extension and 3-pixel fall-off around the entire molecule to produce the final density map with an overall resolution of 3.9 Å for iA2M datasets and 5~8 Å for native datasets. CryoDRGN (6), a deep learning generative neural network, was used to analyze dynamic structures of native A2M and structure variations of iA2M-trypsin. More specifically, for nA2M dataset, a 8 dimensional latent variable model was trained on 257,014 particles (D=150 pixel, 2.08Å per pixel) for 50 epochs. The architecture of encoder and decoder network was set as 1,024×3. After training, automated analyze tools were used for latent space clustering (k=35), particle classification and density map was generated on every cluster center. By using cryoDRGN, two more states in plasma-purified A2M (Fig S2) and two more trypsin-bound states in iA2M (Fig S3) were identified that were not detected by traditional classification. To validate the effectiveness of these new states, particles in each class of cryoDRGN were reconstructed with homogeneous reconstruction in CryoSPARC (7), which reproduced a similar, albeit lower-resolution structure (Figure S9).

**Model building and refinement.** Homologue modeling of iA2M-trypsin was performed in Coot (8), using the crystal structure of human iA2M-MA (PDB ID: 4ACQ) as the initial model. Map refinement was carried out using phenix.real_space_refine (9), with secondary structure and Ramachandran restraints. The structure model of nA2M was built by fitting the structures of sub-domains of iA2M-trypsin into the corresponding densities of nA2M. Specifically, for the density map of nA2M, coordinates of nine macroglobulin-like domains (MG1-MG7, CUB and RBD) and one α-toroid domain (TED) were split from iA2M-trypsin coordinates and fitted into the corresponding sub-domain densities in nA2M monomer map as a rigid-body, which produced a high correlation between each individual domain and the corresponding density. The linkers between domains were then manually built in COOT and refined using phenix.real_space_refine program, with secondary structure and Ramachandran restraints, to finally generate the coordinates of nA2M. Chimera (10), ChimeraX (11) and PyMOL (The PyMOL Molecular Graphics System Version 2.3) were used for graphical visualization.

**Figures and Table**

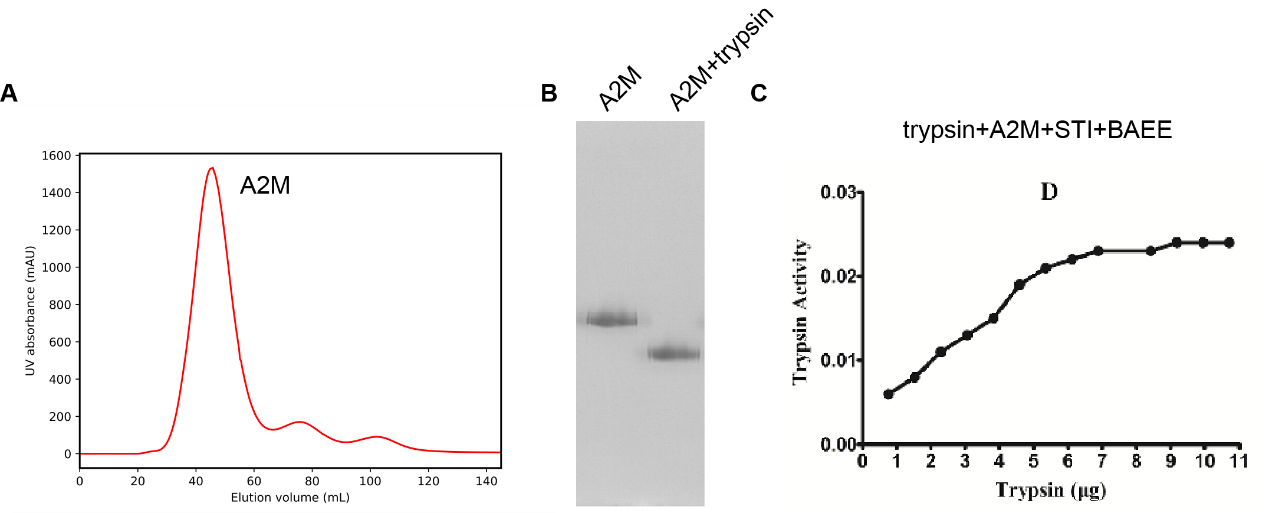

**Fig S1. Purification and characterization of human A2M.**

**(A)** Size-exclusion [chromatography](https://www.baidu.com/link?url=a8FGGjHcZwDazl2wP50AmDZBK_dxfW-qeUcLAgYKBHoKm1dFCrLaxa_Xrd_Y0AodAxEGJngQijUqM1EreU_RkJbgWP5uBNIbKSf8nAFk-1r9Wl12SS4WcX1OieVeJsKwnOFkKfihFU6a_STR2YWqKa&wd=&eqid=ebed989d0021f0f4000000035d4cf502) profile of plasma-purified human A2M. (**B)** A2M (lane1) and iA2M-trypsin (lane2) running in Native PAGE^TM^ 4–16% Bis-Tris gel. A migration was shown for iA2M-trypsin in the native gel. **(C)** Protease entrapment activity of alpha-2-macroglobulin.

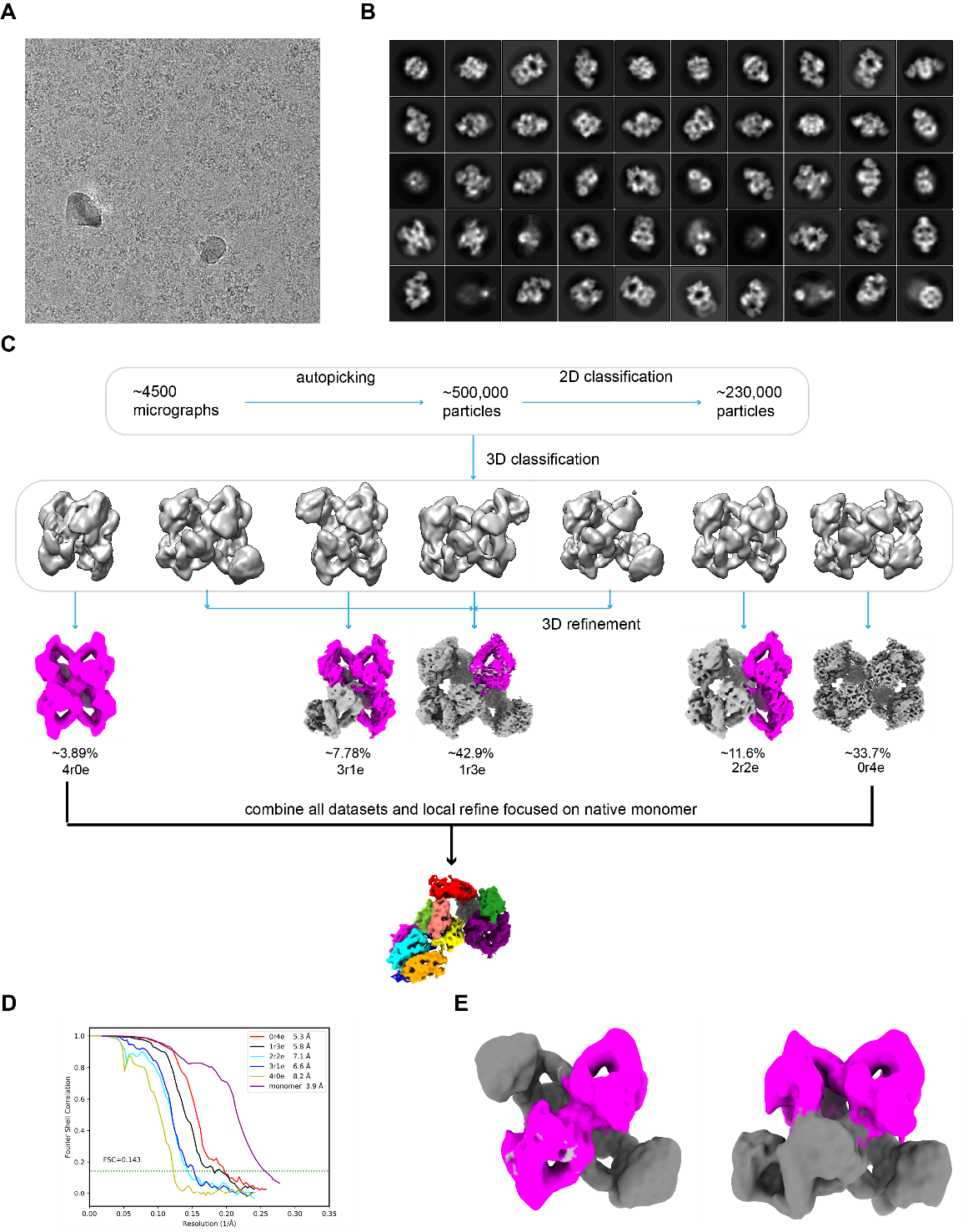

**Fig S2**. **Cryo-EM** **data processing of nA2M.**

**(A-D)** indicate the raw micrograph (A), 2D class averages (B), data processing flowchart (C), and FSC curves (D) of the plasma-purified hA2M reconstruction, respectively. Labels of 0r4e, 1r3e, 2r2e, 3r1e, 4r0e in (C, D) refer to conformations I, II, III, IV, V of hA2M as shown in Fig. 1a respectively, in which “r” represents “retracted” monomer, “e” represents “extended” monomer. (**E**) In addition to the intermediate state of 2r2e showed in (C), two intermediate states consisting of two crossed or horizontal induced monomers were resolved by CryoDRGN in purified hA2M.

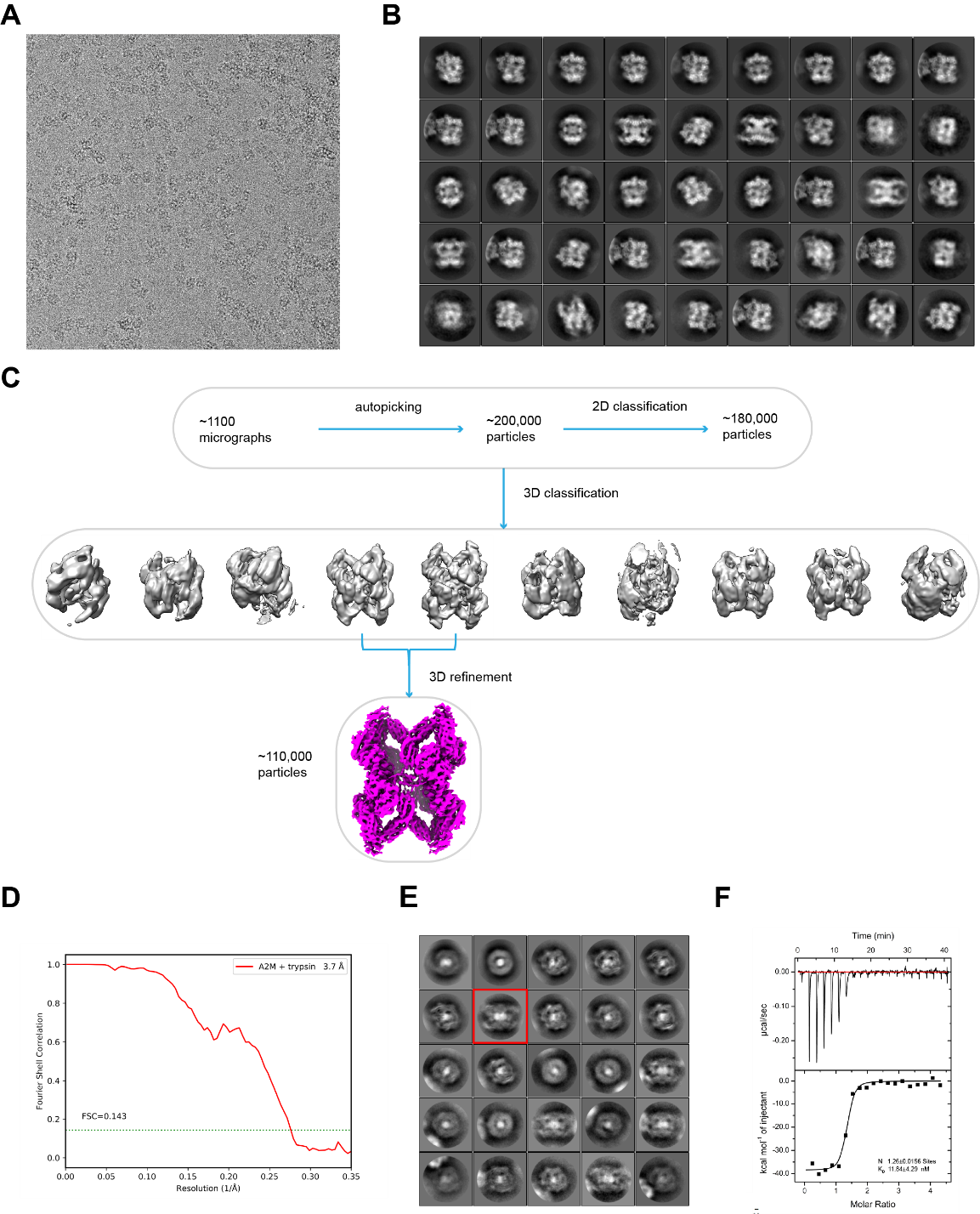

**Fig S3.** **Cryo-EM data processing and ITC profile of iA2M-trypsin.**

**(A-D)** indicate the raw micrograph (A), 2D class averages (B), data processing flowchart (C), and FSC curve (D) of the iA2M-trypsin reconstruction respectively. **(E)** 2D class averages of iA2M-trypsin particles with the outer A2M density subtracted showing the densities of two proteases molecules. **(F)** The [isothermal titration calorimetry](https://www.baidu.com/link?url=w0aIQG_89L8KGzbYLv-4o7zs6O0eOmn8QfOSnVeICpemw4D_eWqrSwTG_7kf5cc1zNu5rPhdK9DxumUbGFcrpufqXUdQfXeuV-k5qqrfeZzDh3oQIE4c8sX6F8ndhJoof3dUotdk266EOkgzYpVLs_&wd=&eqid=e49982d400051c01000000035d5a8917) (ITC) assay of iA2M-trypsin shows the molar ratio of A2M and trypsin. **(G)** Central slices of two intermediate states of iA2M-trypsin resolved by CryoDRGN showing an 1:1 stoichiometric ratio of A2M tetramer with substrate trypsin binding at different locations. **(H)** Reconstructed density of iA2M-trypsin displayed at a low contour level shows the flexible density of RBD (circled).

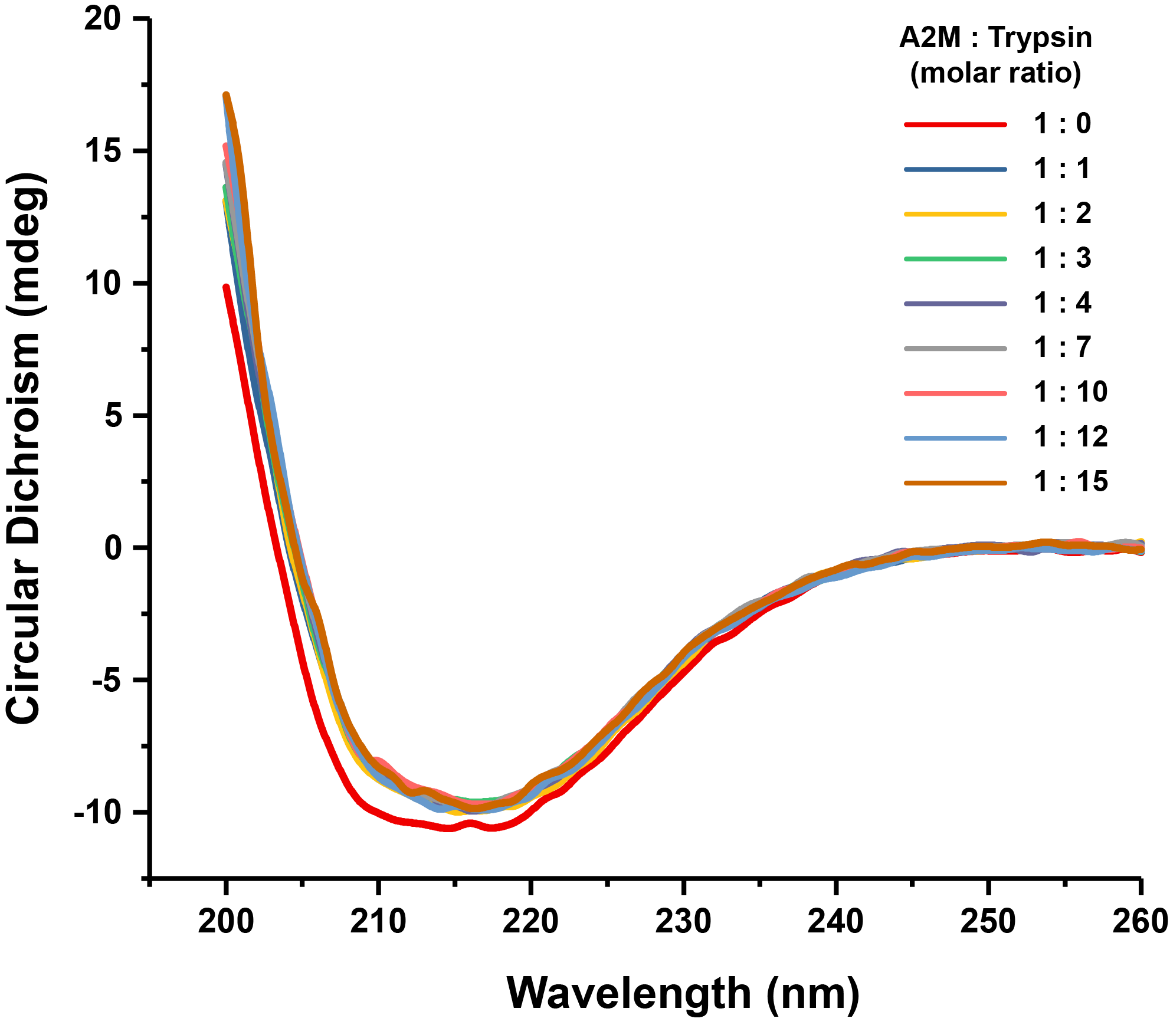

**Fig S4. CD spectra of purified hA2M and iA2M-trypsin.**

CD spectra of purified hA2M and iA2M-trypsin at the A2M:trypsin molar ratio of 1:1, 1:2, 1:3, 1:4, 1:7, 1:10, 1:12, 1:15, respectively, indicating the second structures of A2M were marginally changed when induced by different concentration of proteases.

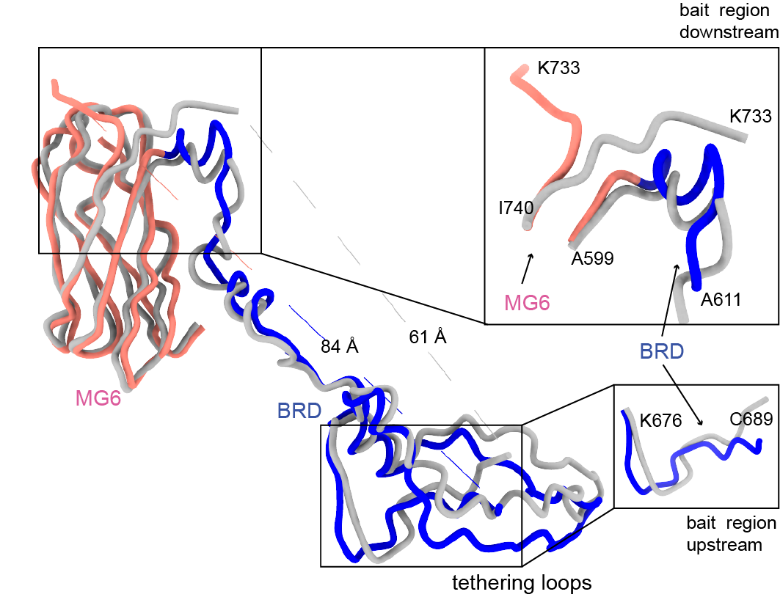

**Fig S5. Structural alignment of MG6 and BRD between nA2M and iA2M-trypsin.**

Structural alignment of MG6 (pink) and BRD (blue) domains between nA2M and iA2M-trypsin was performed using NeeDleman-Wunsch algorithm in UCSF Chimera. Variable regions of native monomer are colored in gray, while rearranged regions of induced monomer are colored as Fig. 2e. The distances of vanished BRD region (689-733) change from 61 Å in native state to 84 Å in induced state. The downstream and upstream of bait region are magnified to show the conformational changes.

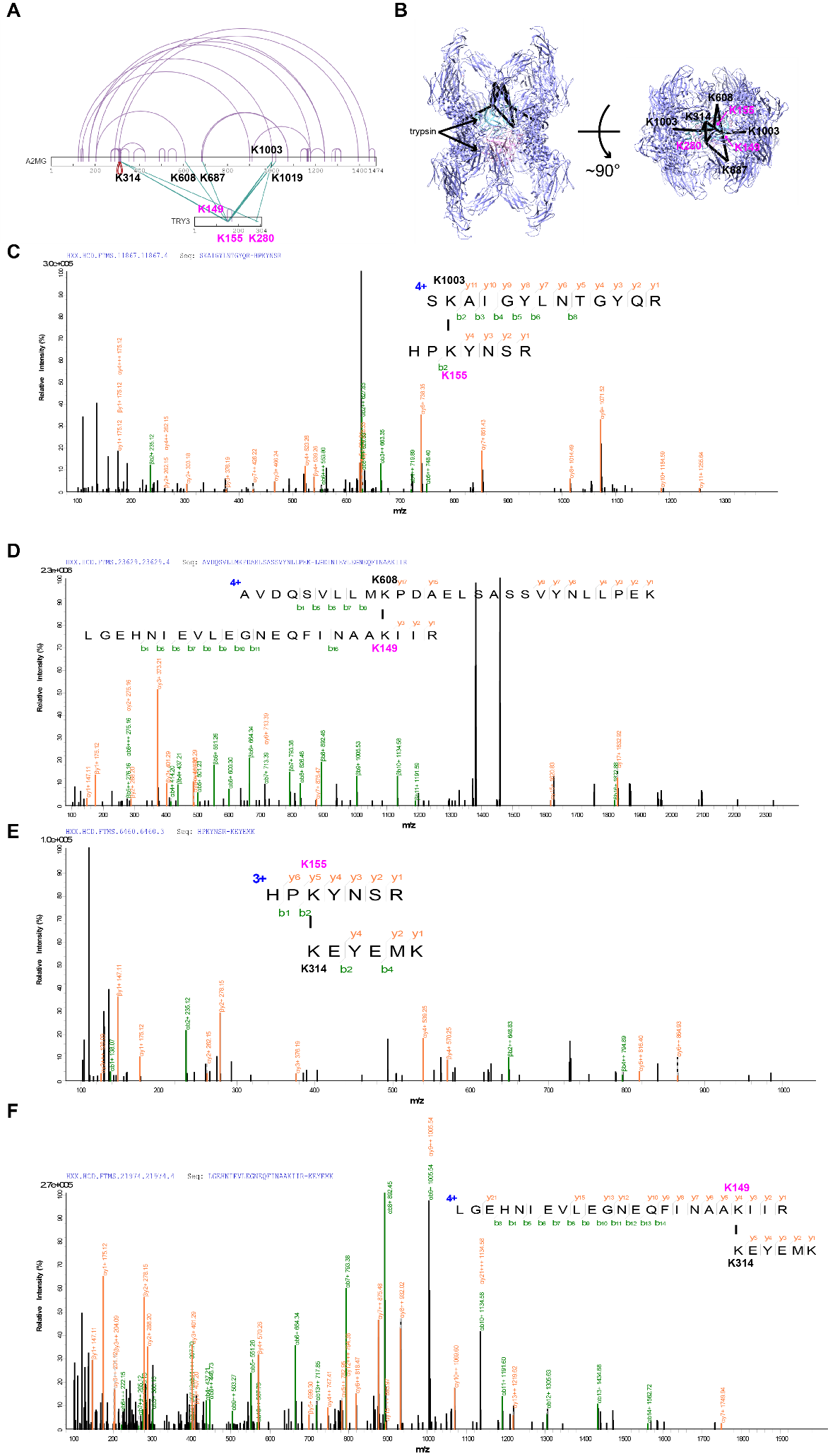

**Fig S6. Cross-link mass spectrum analysis of iA2M-trypsin.**

**(A)** Linkage map showing the identified pairwise cross-links between A2M and trypsin. Critical residues contributing to cross linkage are highlighted. **(B)** Docked structural model of the iA2M-trypsin ternary complex according to CX-MS results. **(C-F)** MS/MS spectrums of pairs of peptides cross-linked by DSS from the iA2M-trypsin complex, including K1003-K155, K608-K149, K314-K155, K314-149 of iA2M-trypsin.

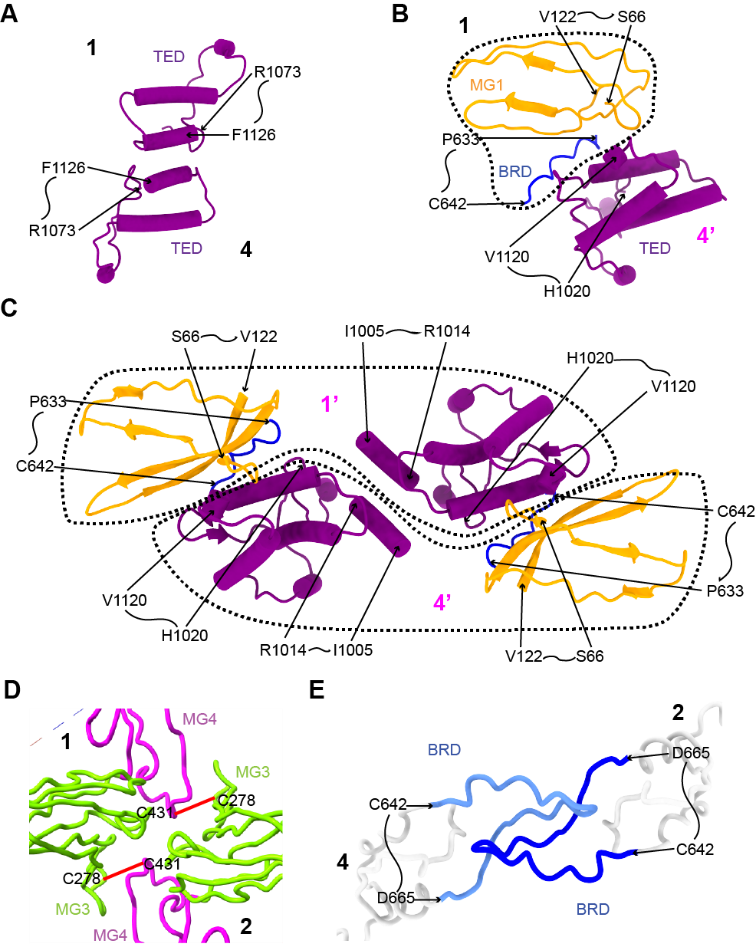

**Fig S7. Magnified views of interactions between subunits of nA2M and iA2M-trypsin as shown in Fig 3.**

**(A-C)** Magnified views of Fig 3 (C-E) showing the interactions between two A2M monomers, including n-n monomers (A), n-i monomers (B), and i-i monomers (C) respectively. **(D)** Magnified view of Fig 3f showing the disulfide bonds (red lines) between the disulfide linked monomers. **(E)** Magnified view of Fig 3G showing the interaction between tethering loops of opposite monomers.

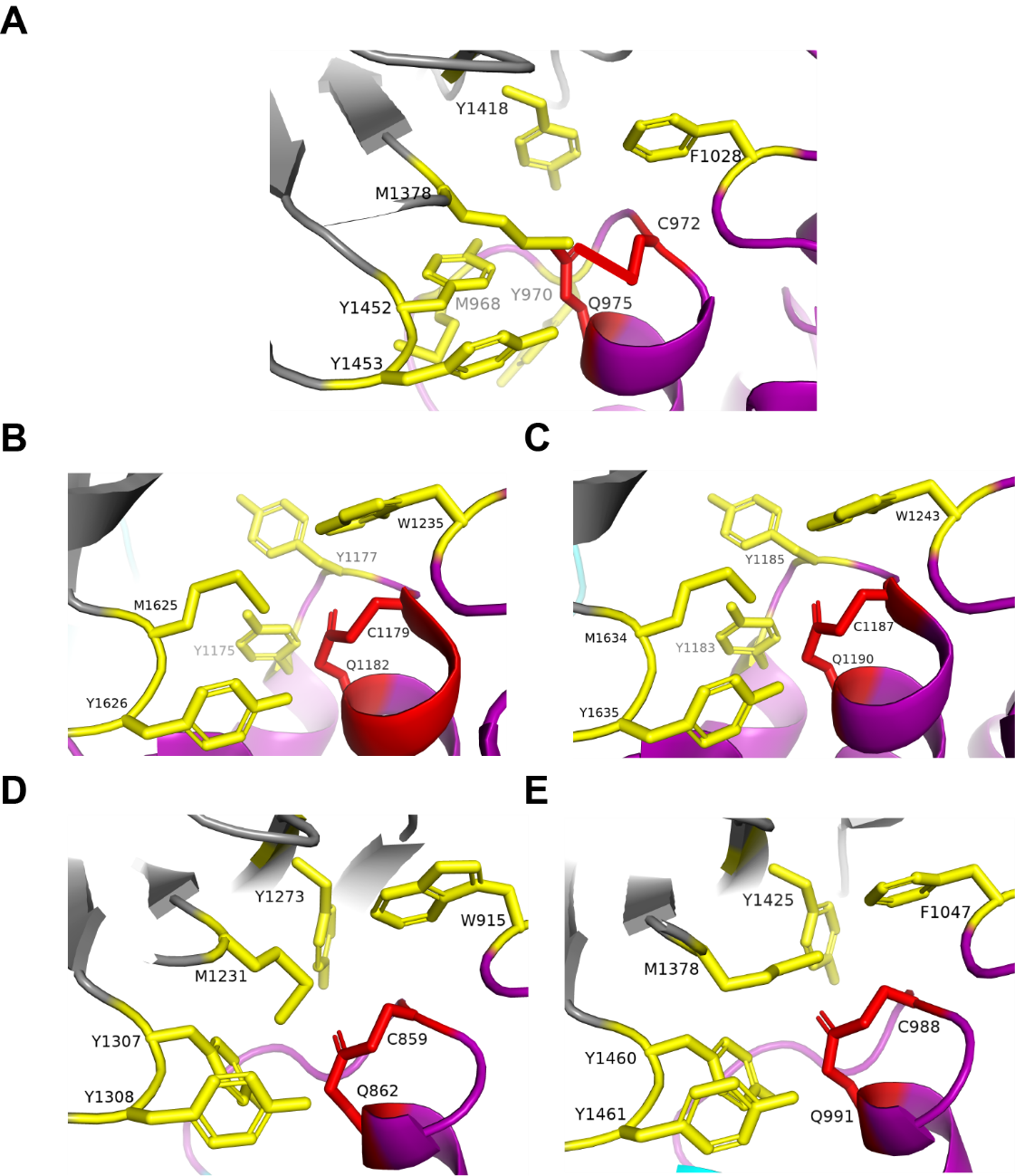

**Fig S8. Key residues contributing to the hydrophobic pocket formation in human A2M and homologues.**

Critical residues constituting the“hydrophobic pocket”in human A2M (A), [prokaryotic](https://www.baidu.com/link?url=frByxZ2psSVV7Ruvbs9rd8AAh2N3bMnUhXoH5o_KzAXcvFSX7axIfYbeIA-3LSkbrMmB_yWMLD0rmPSqCeuI56NM2BhGpDfkchRVMd5QhYC&wd=&eqid=d0aa871900001b53000000035d284919) Salmonella enterica ser A2M (SEAM) (PDB: 4U48) (B), Escherichia coli A2M (ECAM) (PDB: 4ZIU) (C), eukaryotic Anopheles gambiae thiol ester-containing protein TEP1 (PDB: 4D94) (D), and human complement component C3 (PDB: 2A73) (E), are marked respectively. Key residues that contribute to the hydrophobic interactions are colored yellow. Two residues forming the thioester bond are colored red.

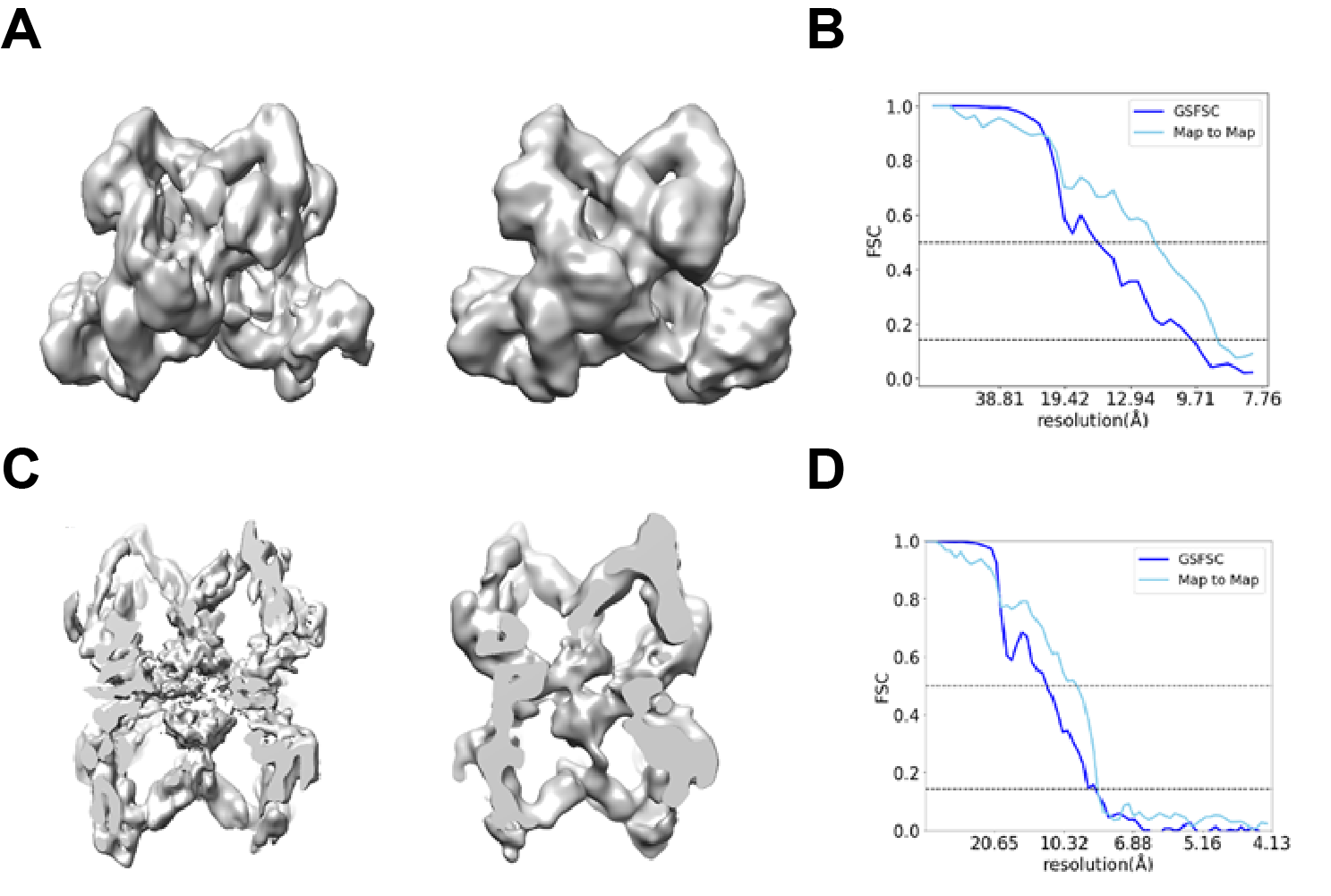

**Fig S9. Validation on the classification results of cryoDRGN.**

**(A)** A representative classification map of nA2M generated by cryoDRGN (left) and the corresponding reconstruction density map by cryoSPARC homogeneous refinement (right) with the same particles. **(B)** Gold standard FSC curve between independent half-maps of the cryoSPARC reconstruction and map-to-map FSC between the cryoDRGN and cryoSPARC maps of nA2M shown in (A). **(C)** A representative classification map of iA2M-trypsin generated by cryoDRGN (left) and the corresponding reconstruction density map by cryoSPARC homogeneous refinement (right) with the same particles. **(D)** Gold standard FSC curve between independent half-maps of the cryoSPARC reconstruction and map-to-map FSC between the cryoDRGN and cryoSPARC maps of iA2M shown in (C).

**Table S1. Data collections and refinement statistics of the purified A2M and iA2M-trypsin.**

| **Dataset** | **Purified A2M (1r3e)** | **iA2M-trypsin** |
| --- | --- | --- |
| **Data collection** |  |  |
| EM equipment | FEI Titan Krios | FEI Titan Krios |
| Voltage (kV) | 300 | 300 |
| Detector | K2 Summit | K2 Summit |
| Pixel size (Å) | 1.04 | 1.04 |
| Electron dose (e^–^/Å^2^) | 60 | 60 |
| Defocus range (μm) | -1.8~-2.5 | -1.8~-2.5 |
| **Reconstruction** |  |  |
| Software | RELION 3.0 + cisTEM + cryoSPARC + cryoDRGN | RELION 3.0 + cisTEM + cryoSPARC+ cryoDRGN |
| Particle Numbers | 98,600 | 110,000 |
| Symmetry | C1 | D2 |
| FSC threshold | 0.143 | 0.143 |
| Final resolution (Å) | 5.2 | 3.9 |
| Map-sharpening *B* factor (Å^2^) | -230 | -185.2 |
| **Model building** |  |  |
| Software | Coot | Coot |
| **Refinement** |  |  |
| Software | Phenix | Phenix |
| CC_mask | 0.7197 | 0.7898 |
| Bond lengths (Å) | 0.012 | 0.009 |
| Bond angles (°) | 1.427 | 1.262 |
| **Validation** |  |  |
| Clashscore | 18.33 | 9.87 |
| Rotamers outliers (%) | 1.31 | 1.25 |
| **Ramachandran plot** |  |  |
| Favored (%) | 85.72 | 89.25 |
| Allowed (%) | 13.63 | 10.02 |
| Outliers (%) | 0.65 | 0.73 |

**Movie S1. Conformational rearrangements from native A2M to induced A2M.**
